# A common neural substrate for elevated PTSD symptoms and reduced pulse rate variability in combat-exposed veterans

**DOI:** 10.1101/364455

**Authors:** Daniel W. Grupe, Ted Imhoff-Smith, Joseph Wielgosz, Jack B. Nitschke, Richard J. Davidson

## Abstract

**Background:** Previous studies have identified reduced heart rate variability (HRV) in posttraumatic stress disorder (PTSD), which may temporally precede the onset of the disorder. A separate line of functional neuroimaging research suggests that the ventromedial prefrontal cortex (vmPFC) — a key aspect of a descending neuromodulatory system that exerts inhibitory control over heart rate — shows functional and structural abnormalities in PTSD. No research to date, however, has simultaneously investigated whether altered vmPFC activation is associated with reduced HRV and elevated PTSD symptoms in the same individuals.

**Methods:** We collected fMRI data during alternating conditions of threat of shock and safety from shock in 51 male, combat-exposed veterans with either high or low levels of PTSD symptoms. Pulse rate variability (PRV) – an HRV surrogate calculated from pulse oximetry – was assessed during a subsequent resting scan. Correlational analyses tested for hypothesized relationships between vmPFC activation, PRV, and distinct dimensions of PTSD symptomatology.

**Results:** Re-experiencing PTSD symptoms were inversely associated with high-frequency (HF)-PRV, thought to primarily reflect parasympathetic control of heart rate, in veterans with elevated PTSD symptoms. Lower HF-PRV was associated with reduced vmPFC activation for the contrast of safety-threat in a region that also showed an inverse relationship with re-experiencing symptoms.

**Conclusions:** Reduced vmPFC responses to safety vs. threat were associated with both reduced HF-PRV and increased re-experiencing symptoms. These results tie together previous observations of reduced HRV/PRV and impaired vmPFC function in PTSD and call for further research on reciprocal brain-body relationships in understanding PTSD pathophysiology.

## 1. Introduction

Trauma is an embodied experience. In addition to the psychological consequences of trauma, somatic symptoms cause extreme distress in posttraumatic stress disorder (PTSD) and pose a substantial hurdle in the treatment of and recovery from trauma (van der Kolk, 2014). In spite of this, etiological theories of PTSD – such as dominant fear learning perspectives – typically emphasize neural mechanisms and treat somatic symptoms as epiphenomenal; consequently, first-line behavioral interventions primarily target psychological processes. The investigation of relationships between the brain and periphery, and how these relationships are altered in trauma-exposed individuals, may elucidate novel candidate mechanisms of PTSD etiology. These candidate mechanisms can then be used as treatment targets for somatically focused, “alternative” treatments for PTSD (Gallegos, Crean, Pigeon, & Heffner, 2017; Polusny et al., 2015) that may be better tolerated or more effective for some traumatized individuals.

One way to learn more about such mechanisms is by assessing peripheral psychophysiology in conjunction with brain imaging techniques. A peripheral index of particular interest in this regard is heart rate variability (HRV), which is thought to reflect adaptive regulatory control of autonomic function by central mechanisms (Thayer & Lane, 2000). Of particular interest is high-frequency HRV, the parasympathetically dominated component of HRV tied to the respiration cycle (typically in the frequency of 0.12–0.40 or 0.15–0.40 Hz; Allen, Chambers, & Towers, 2007). Consistent with a role for HRV in flexible regulatory control, reduced HRV is seen in psychiatric disorders marked by deficient inhibitory control of emotional and physiological responding, including depression (Kemp et al., 2010) and anxiety disorders (Chalmers, Quintana, Abbott, & Kemp, 2014). A meta-analysis of 19 studies comparing PTSD patients to controls demonstrated reduced HRV in PTSD, particularly for the parasympathetically dominant HF-HRV (Nagpal, Gleichauf, & Ginsberg, 2013). Further, two large studies in pre-deployment soldiers found that reduced HF-HRV prior to combat exposure (or, similarly, a smaller HF/low-frequency ratio) predicted post-deployment PTSD symptoms (Minassian et al., 2015; Pyne et al., 2016).

Neurally, increased heart rate variability is associated with increased activation of the ventromedial prefrontal cortex (vmPFC), as demonstrated most clearly in a meta-analysis of neuroimaging studies by Thayer and colleagues (2012). The authors suggested that descending projections from the vmPFC and other aspects of a descending “visceromotor system” provide a critical regulatory role over the autonomic nervous system and support context-appropriate threat responding. Notably, the vmPFC has also been extensively implicated in the pathophysiology of PTSD, as underscored by quantitative meta-analyses of functional neuroimaging research comparing patients and controls (Hayes, Hayes, & Mikedis, 2012; Stark et al., 2015). In particular, reduced activation of the vmPFC during extinction recall or in response to cues representing safety has been noted in PTSD (Garfinkel et al., 2014; Grupe, Wielgosz, Davidson, & Nitschke, 2016; Milad et al., 2009).

Collectively, these data suggest a mechanistic link between lower vmPFC activity and reduced HRV, which may account for multiple aspects of PTSD symptomatology in trauma-exposed individuals. Because HRV is proposed to index physiological flexibility in response to changing environmental demands, vmPFC dysfunction and consequently reduced HRV could compromise traumatized individuals’ ability to respond adaptively across dynamic contexts, resulting in contextually inappropriate and overgeneralized threat responding best encapsulated by hypervigilance and other symptoms of hyperarousal. Alternatively, reduced regulatory control reflected in lower levels of vmPFC activity and reduced HRV may allow unwanted traumatic memories or flashbacks to emerge (Gillie & Thayer, 2014), consistent with a broader proposed role linking lower HRV to reduced inhibitory control of thoughts, emotions, and physiology in a broad array of anxiety disorders (Chambers et al., 2014). However, the hypothesis that compromised vmPFC function and reduced HRV reflect a common mechanism in the pathology of PTSD remains somewhat theoretical, as few if any studies have simultaneously explored relationships among PTSD symptoms, vmPFC function, and HRV in the same participants.

In the current study, we investigated each of these factors in a functional MRI study of combat trauma-exposed veterans. Participants took part in an fMRI task involving alternating conditions of unpredictable threat and safety. We collected pulse-rate data using pulse oximetry during a subsequent resting-state scan owing to the difficulty of collecting electrocardiography data in the MRI environment, and analyzed high-frequency pulse rate variability, or HF-PRV. Importantly, PRV and HRV, which capture different physiological readouts of cardiac variability, are highly correlated during resting conditions (Hayano, Barros, Kamiya, Ohte, & Yasuma, 2005; Schäfer & Vagedes, 2013). We first tested the hypothesis that elevated PTSD symptoms would be associated with reduced resting HF-PRV. Follow-up analyses revealed that this hypothesized relationship was most prominent for re-experiencing symptoms of PTSD. Second, we hypothesized that reduced vmPFC activation during safety from shock would be associated with lower resting HF-PRV. We identified this relationship in a sector of the vmPFC (BA10) in which we previously demonstrated a correlation between reduced activation and elevated re-experiencing symptoms (Grupe et al., 2016). Our results suggest a common neural substrate for reduced PRV and elevated re-experiencing symptoms in the same combat-exposed veterans, providing preliminary support for a hypothesized brain-body mechanism that could serve as a novel treatment target in future intervention studies.

## 2. Methods

### 2.1 Participants

Participants for this study were veterans of Operation Enduring Freedom/Operation Iraqi Freedom (OEF/OIF) who were exposed to one or more life-threatening war zone trauma events during deployment. These individuals were recruited through online and community advertisements and in collaboration with the Madison VA Hospital, the Madison Veterans’ Center, the Wisconsin National Guard, and other veterans’ organizations. We previously reported on relationships between individual PTSD symptom clusters and brain responses to unpredictable threat anticipation in this sample (Grupe et al., 2016).

Following written informed consent, a team of clinically trained interviewers administered the Clinician-Administered PTSD Scale (CAPS; Blake et al., 1990) and Structured Clinical Interview for DSM-IV (SCID; First et al., 2002) with supervision from a licensed clinical psychologist (JBN). Exclusionary conditions included substance dependence within the past 3 months and current or past bipolar, psychotic, or cognitive disorders. Based on CAPS scores, individuals were enrolled into a combat-exposed control (CEC) group or a posttraumatic stress symptoms (PTSS) group. Members of the CEC group were free of current Axis I disorders and had CAPS scores < 10, and members of the PTSS group had PTSD symptoms occurring at least monthly with moderate intensity and CAPS scores ≥ 20. Current major depression or dysthymia was not exclusionary in the PTSS group. Current treatment with psychotropic medications (other than benzodiazepines or beta-blockers) or maintenance psychotherapy was permitted if treatment was stable for 8 weeks prior to the beginning of the study (see Supplementary Table 1 for complete participant characteristics).

Although 58 participants were enrolled, we only analyzed data from male participants as the sample included only 4 women. In addition, 2 participants could not tolerate the shock and 1 exceeded the motion threshold during fMRI scanning. The final sample of 51 male veterans consisted of 17 in the CEC group and 34 in the PTSS group, 16 of whom met full PTSD diagnostics criteria and 18 of whom met criteria for only 1 or 2 of the symptom clusters (**Table 1**).

**Table 1:**
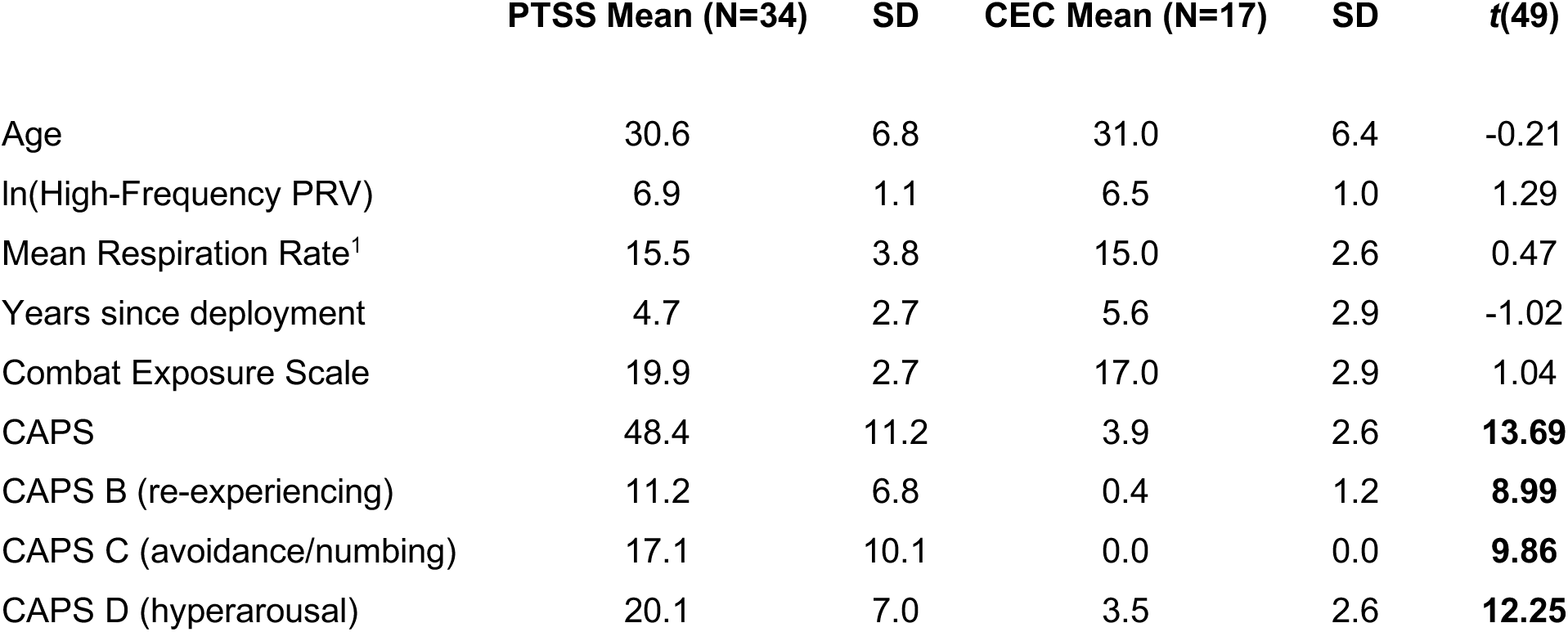
Sample characteristics. Demographic and clinical variables for all participants and results of independent samples *t* tests comparing participants in the posttraumatic stress symptoms (PTSS) and combat exposed control (CEC) groups. **Notes**: bolded values = group differences at *p* < 0.05. PRV = pulse rate variability; CAPS = Clinician Administered PTSD Scale. ^1^Only 44 participants had valid respiration data.

|  | PTSS Mean (N=34) | SD | CEC Mean (N=17) | SD | *t*(49) |
| --- | --- | --- | --- | --- | --- |
| Age | 30.6 | 6.8 | 31.0 | 6.4 | -0.21 |
| ln(High-Frequency PRV) | 6.9 | 1.1 | 6.5 | 1.0 | 1.29 |
| Mean Respiration Rate<sup>1</sup> | 15.5 | 3.8 | 15.0 | 2.6 | 0.47 |
| Years since deployment | 4.7 | 2.7 | 5.6 | 2.9 | -1.02 |
| Combat Exposure Scale | 19.9 | 2.7 | 17.0 | 2.9 | 1.04 |
| CAPS | 48.4 | 11.2 | 3.9 | 2.6 | **13.69** |
| CAPS B (re-experiencing) | 11.2 | 6.8 | 0.4 | 1.2 | **8.99** |
| CAPS C (avoidance/numbing) | 17.1 | 10.1 | 0.0 | 0.0 | **9.86** |
| CAPS D (hyperarousal) | 20.1 | 7.0 | 3.5 | 2.6 | **12.25** |

### 2.2 fMRI task

We previously published complete details of the fMRI task in this sample (Grupe et al., 2016). Briefly, during a baseline visit within 2 weeks of the MRI scan, participants took part in a shock calibration procedure to determine a level of shock, delivered to the right ventral wrist, that was perceived as “very unpleasant, but not painful”. On the day of the MRI scan a single shock was delivered to confirm the shock calibration procedure, and participants took part in an instructed threat anticipation task (**Figure 1A**). Each trial began with a 2-s presentation of a blue or yellow square, indicating threat of shock or safety from shock (counterbalanced). Next, the same color clock appeared for 4–10 s (mean duration = 7.67 s). On predictable trials, a red mark appeared in a random location and the anticipation period ended when a slowly rotating hand reached this mark. On unpredictable trials, no red mark appeared and participants could not predict the end of the anticipation period. The current analyses focused on unpredictable trials, as in Grupe et al., (2016). On 12/42 threat trials, a 200-ms electric shock was delivered concurrently with a neutral tone. On all other trials, the anticipation period concluded with a 200-ms tone only. All trials were followed by a 5–9 s inter-trial interval. The scan session included 42 threat trials and 30 safe trials, split evenly between predictable/unpredictable conditions, allowing us to analyze an equivalent number (15 each) of non-reinforced unpredictable threat and safe trials. For the current study, we analyzed data from the unpredictable conditions only, contrasting functional activation during the 4–10s anticipation epoch between unpredictable threat (uThreat) and unpredictable safe (uSafe) trials.

**Figure 1.**
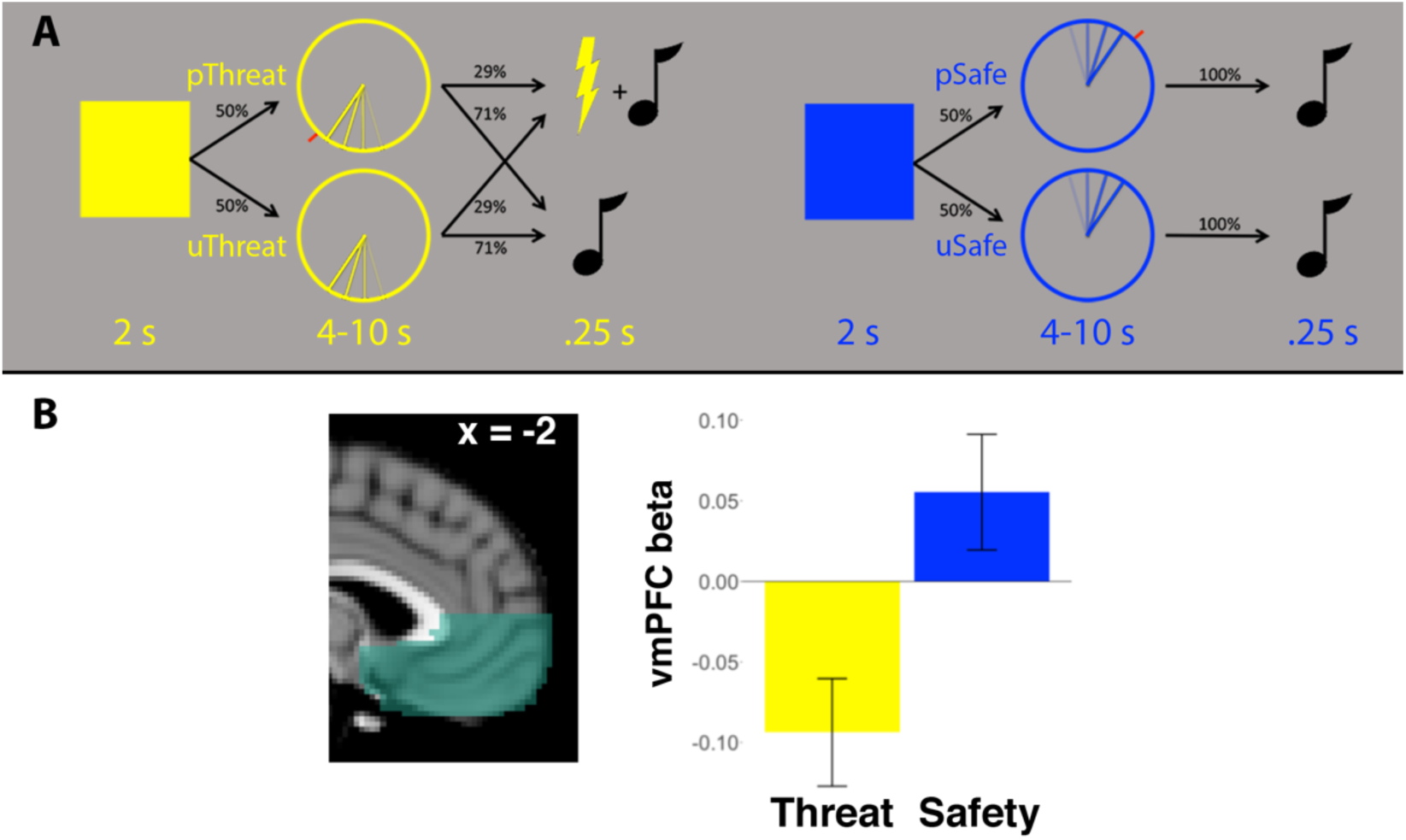
fMRI task. (A) During the threat anticipation task, the color of a 2s square indicated threat of shock or safety from shock (counterbalanced). A subsequent 4–10s anticipation period terminated in shock on 29% of trials in the threat condition. Analyses focused on unpredictable trials (bottom row), where participants had no visual cue to indicate the termination of the anticipation period. (B) Across the anatomically defined ventromedial prefrontal cortex (vmPFC) ROI, the group as a whole showed greater activation during safe vs. threat trials (*t*(50) = 6.29, *p* < 0.001).

### 2.3 MRI data collection

MRI data were collected on a 3T X750 GE Discovery scanner using an 8-channel head coil and ASSET parallel imaging with an acceleration factor of 2. Data collected included 3 sets of echo planar images (EPIs) during the threat anticipation task (240 volumes/8:00, TR=2000, TE=20, flip angle=60°, field of view=220 mm, 96×64 matrix, 3-mm slice thickness with 1-mm gap, 40 interleaved sagittal slices), 1 set of EPIs during a subsequent resting-state scan (210 volumes/7:00) and a T1-weighted anatomical image for functional data registration. Visual stimuli were presented using Avotec fiberoptic goggles, auditory stimuli were presented binaurally using Avotec headphones, and behavioral responses were recorded using a Current Designs button box.

### 2.4 Psychophysiology data collection and processing

Peripheral physiological data were acquired during each of the 3 task runs and the subsequent resting-state scan; analyses here utilized data from the 7-minute resting-state scan. Pulse rate data were acquired using a pulse oximeter on the second finger of the left hand (contralateral to shock delivery), and respiration data were acquired using a belt placed at the bottom of the ribcage. All peripheral physiological data were amplified using a BIOPAC MP-150 system and digitized at 1000 Hz.

Pulse rate data were preprocessed using in-house Matlab software that automatically detected heartbeats, after which missing or extra beats were manually identified. The resulting time series of interbeat intervals (IBIs) was analyzed in CMetX (Allen et al., 2007). Detected artifacts (>300 ms difference between consecutive IBIs) were manually reviewed and rejected if determined to be artifactual. The primary outcome of interest was high-frequency pulse rate variability (HF-PRV; (Schäfer & Vagedes, 2013)), defined as the natural log of the band-pass filtered IBI time series between 0.12–0.40 Hz, or the typical frequency range for the respiratory cycle.

Mean respiration rate was calculated using in-house Matlab scripts. We visually inspected the respiration signal and filtered out distorted or artifactual data before extracting trough-to-trough intervals from useable data and calculating mean respiration rate (breaths/minute). We excluded participants from respiration analyses if they had less than 60s of clean data from which we could estimate respiration rate.

### 2.5 Data analysis

Statistical analyses were conducted in R version 3.2.2. All fMRI processing and analysis was conducted using FEAT (FMRI Expert Analysis Tool) Version 6.00, part of FSL (FMRIB’s Software Library, www.fmrib.ox.ac.uk/fsl), as fully described in our previous publication (Grupe et al., 2016).

To test hypothesis 1 (inverse relationship between PRV and PTSD symptoms), we first conducted an independent samples *t* test to compare HF-PRV between PTSS and CEC groups. We also calculated Pearson correlations between HF-PRV and total CAPS scores, both in the full sample (N=51) and the PTSS group (N=34). In follow-up analyses we tested whether a significant inverse correlation identified in the PTSS group was specific to individual symptom clusters (re-experiencing, avoidance/numbing, and hyperarousal), first by calculating Pearson correlations between each of the 3 symptom clusters, and second by simultaneous regression of HF-PRV on these 3 clusters. Because HF-PRV is coupled with respiration, we re-ran these analyses using residualized HF-PRV after regressing out resting respiration rate (due to poor respiration data quality in some participants, we could only estimate respiration rate in 29/34 PTSS and 15/17 CEC participants).

To test hypothesis 2, we conducted voxelwise correlation analysis of HF-PRV and uSafe-uThreat contrast estimates constrained to the anatomically defined vmPFC, in the full sample and the PTSS group alone. We again ran follow-up analyses using respiration-residualized HF-PRV. Repeating an analysis from a previous report in this sample linking vmPFC activation to re-experiencing symptoms of PTSD (Grupe et al., 2016), we also conducted simultaneous voxelwise regression of vmPFC contrast estimates on each of the 3 PTSD symptom clusters. The vmPFC mask consisted of medial portions of Brodmann Areas 10, 11, 12, 24, 25, and 32 ventral to the genu of the corpus callosum (generated using the Wake Forest University PickAtlas; Maldjian et al., 2003). Exploratory regression analyses were conducted across the entire brain. Cluster threshold correction was applied to the vmPFC and across the whole brain using a voxelwise threshold of *p* < 0.005, resulting in corrected significance of *p* < 0.05.

## 3. Results

### 3.1 Pulse rate variability is inversely associated with PTSD re-experiencing symptoms

There was no significant difference in HF-PRV between participants in the PTSS and CEC groups (*t*(49) = 1.29, *p* = 0.20, *d* = 0.38), nor was there a relationship between HF-PRV and PTSD CAPS symptoms measured continuously within the full sample (*r*(49) = −0.05, 95% CI = [−0.32, 0.23], *p* = 0.72).

Within the PTSS group alone, there was a significant inverse relationship between HF-PRV and total CAPS symptoms (*r*(32) = −0.41, 95% CI = [−0.66, −0.08], *p* = 0.015; **Figure 2a**). This relationship was strongest for the re-experiencing cluster (*r*(32) = −0.48, 95% CI = [−0.70, −0.17], *p* = 0.004; **Figure 2b**), and was not significant for avoidance/numbing (*r*(32) = −0.31, 95% CI = [−0.59, 0.03], *p* = 0.08) or hyperarousal symptoms (*r*(32) = −0.19, 95% CI = [−0.50, 0.16], *p* = 0.28). To test the specificity of this relationship to re-experiencing symptoms, simultaneous regression of HF-PRV on these 3 symptom clusters was conducted. This revealed a significant relationship between re-experiencing symptoms and HF-PRV (*t*(30) = −2.39, *p* = 0.023), and no relationships for avoidance/numbing or hyperarousal symptoms (*t*s < 1, *p*s > 0.4).

**Figure 2.**
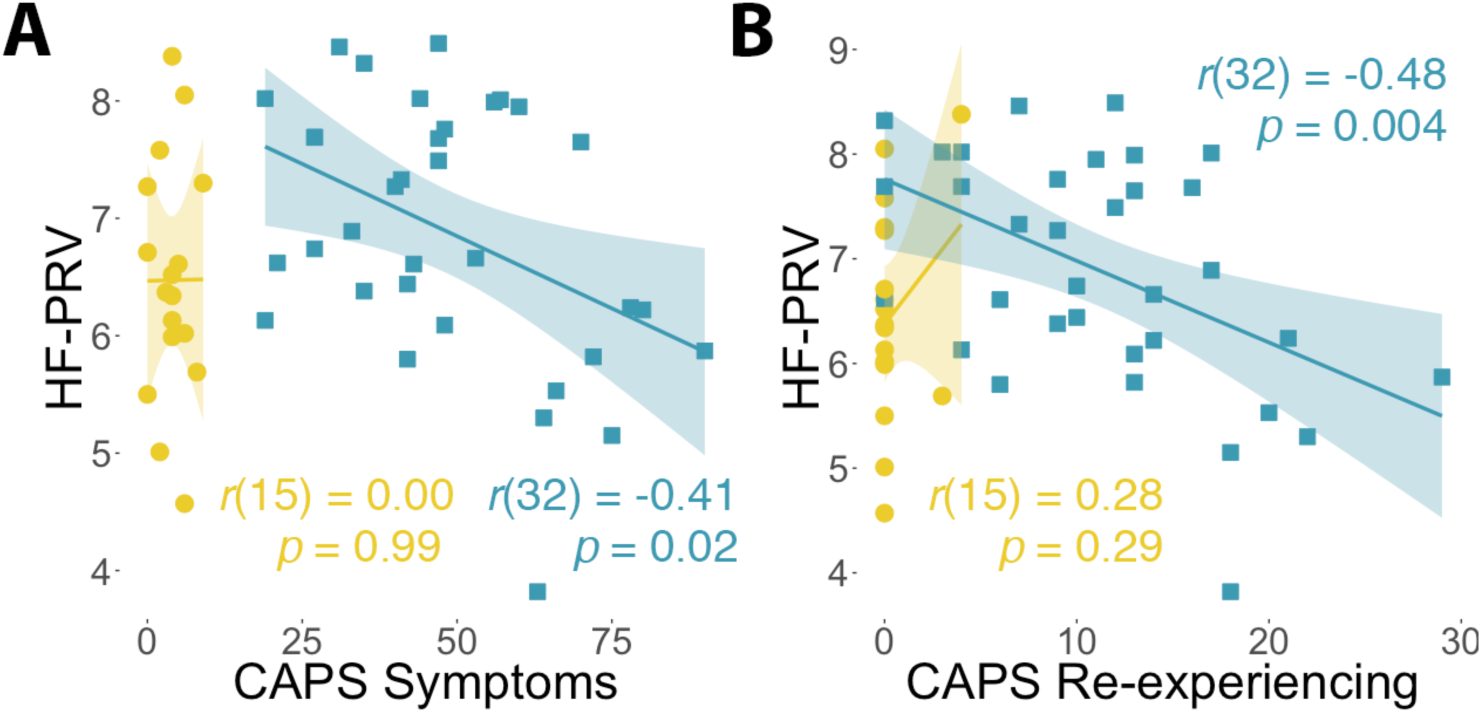
PTSD symptoms and pulse rate variability. (A) Across the entire sample, there was no correlation between total PTSD symptom severity on the Clinician-Administered PTSD Scale (CAPS) and high-frequency pulse rate variability (HF-PRV; *r*(49) = −0.04, *p* = 0.72). There was also no correlation in the control group (yellow), but there was a significant inverse correlation in the posttraumatic stress symptoms (PTSS) group (blue). (B) This inverse relationship in the PTSS group was strongest for re-experiencing CAPS symptoms.

We conducted additional analyses to ensure that individual differences in respiration rate were not driving relationships between HF-PRV and re-experiencing symptoms. Of the 44 participants with valid respiration data, 43 had respiration frequencies within the 0.12–0.40 Hz window used to band-pass filter the IBI time series (the 44^th^ participant fell just outside this window, frequency=0.44 Hz). There was a non-significant inverse relationship between respiration rate and HF-PRV, such that faster breathing tended to be associated with lower HF-PRV (full sample *r*(42) = 0.24, 95% CI = [−0.06, 0.50], *p* = 0.12; PTSS *r*(27) = 0.29, 95% CI = [−0.09, 0.59], *p* = 0.12). Importantly, however, respiration rate was unrelated to overall CAPS symptoms (full sample *r*(42) = 0.19, 95% CI = [−0.11, 0.46], *p* = 0.22; PTSS *r*(27) = 0.24, 95% CI = [−0.14, 0.56], *p* = 0.20) or re-experiencing symptoms (full sample *r*(42) = 0.17, 95% CI = [−0.13, 0.44], *p* = 0.27; PTSS *r*(27) = 0.17, 95% CI = [−0.21, 0.51], *p* = 0.37). Additionally, we also observed a significant correlation in the PTSS group between respiration-adjusted HF-PRV and re-experiencing symptoms (*r*(27) = −0.43, 95% CI = [−0.69, −0.08], *p* = 0.02). In the simultaneous regression model with all 3 symptom clusters, re-experiencing symptoms no longer accounted for significant unique variance in respiration-adjusted HF-PRV (*t*(25) = −1.89, *p* = 0.07).

### 3.2 Reduced PRV and elevated re-experiencing symptoms are associated with reduced activation of a common vmPFC region

Voxelwise regression of uSafe-uThreat contrast estimates revealed small volume-corrected clusters within the vmPFC that showed an inverse relationship with HF-PRV for the full sample (**Figure 3a**) and the PTSS group **Figure 3b**). Participants with lower HF-PRV showed relatively less anticipatory activation for safety vs. threat (greater safe vs. threat activation being the normative pattern across the vmPFC; **Figure 1b**). Each of these clusters was localized to BA10, or the medial frontopolar aspect of vmPFC. Similar small volume-corrected clusters were observed for respiration-adjusted HF-PRV in participants with valid respiration data. Outside of the vmPFC, we identified a whole-brain significant cluster in left middle frontal gyrus that showed a similar inverse relationship with HF-PRV within the PTSS group (**Figure 3c**). No clusters survived whole-brain significance for the full sample.

**Figure 3.**
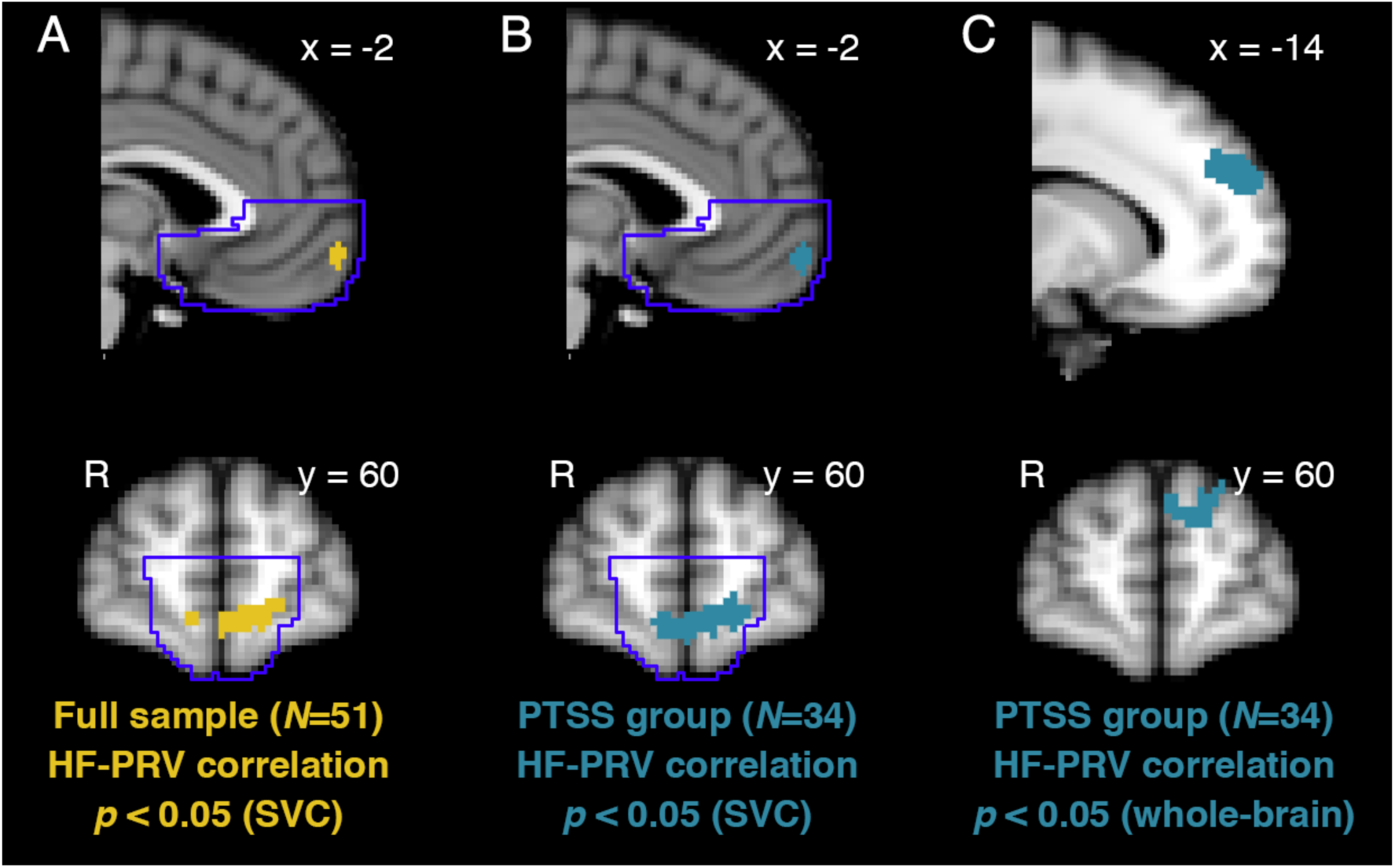
Pulse rate variability and brain responses to safety. (A) Across the entire sample, there was a positive correlation between high-frequency pulse rate variability (HF-PRV) and BOLD responses to safety vs. threat anticipation within the small-volume corrected (SVC) ventromedial prefrontal cortex (vmPFC), corresponding to BA10. (B) This relationship was also observed in the posttraumatic stress symptoms (PTSS) group alone. (C) Outside of the vmPFC, the PTSS group also showed a positive correlation between HF-PRV and BOLD responses to safety vs. threat in the left middle frontal gyrus.

In our previous report on this sample, we identified an inverse relationship in a similar aspect of BA10 between uSafe-uThreat activation and symptoms of re-experiencing (Grupe et al., 2016, Figure S5b). Overlaying statistical maps from the HF-PRV and re-experiencing regression analyses, we identified an overlapping set of voxels (208 mm^3^ volume) in which uSafe-uThreat activation was positively correlated with HF-PRV and negatively correlated with re-experiencing symptoms (**Figure 4**).

**Figure 4.**
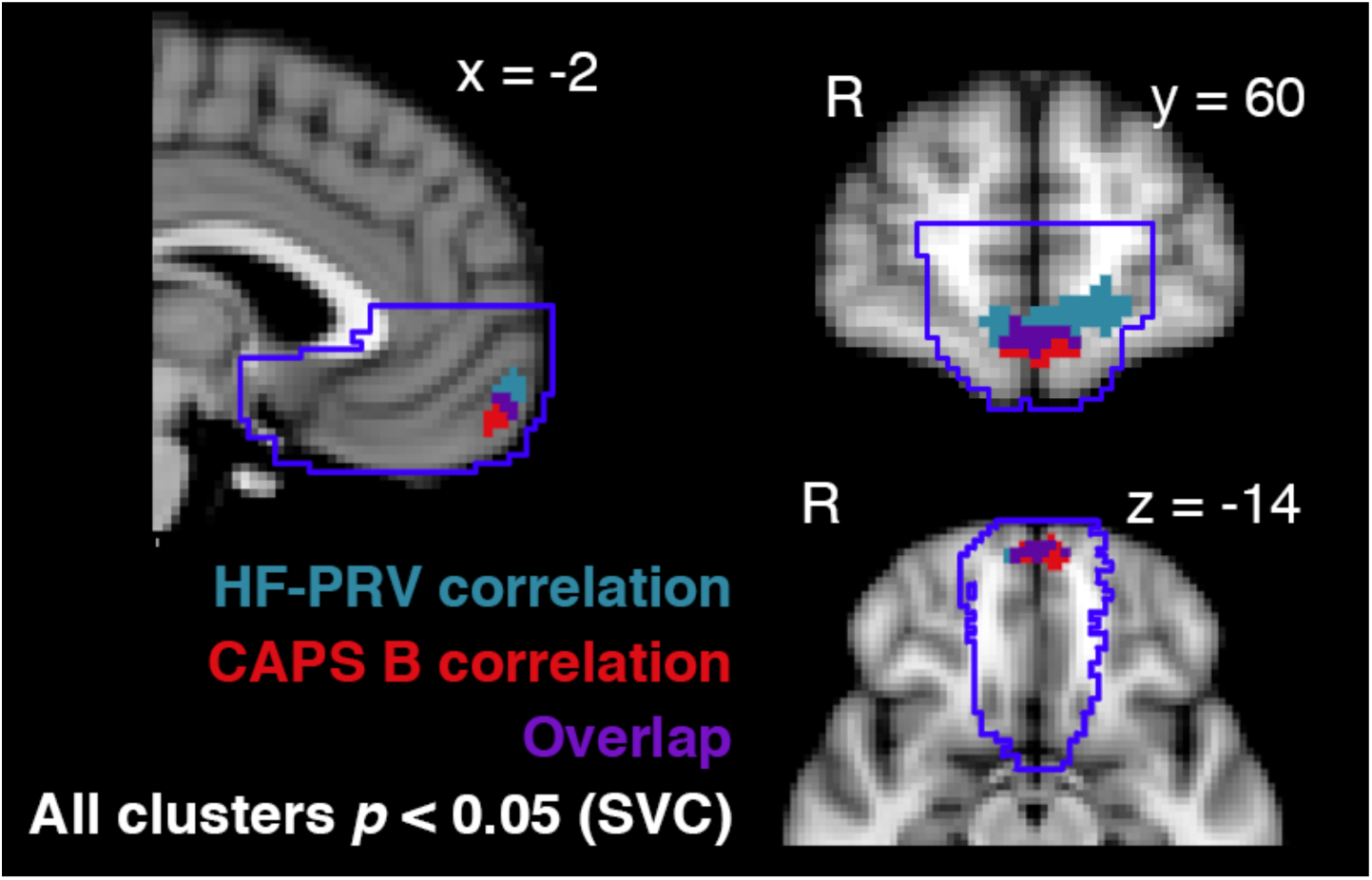
Overlapping vmPFC region for pulse rate variability and re-experiencing symptoms. Within the posttraumatic stress symptoms (PTSS) group, BOLD responses to safety vs. threat were correlated with both high-frequency pulse rate variability (HF-PRV; blue) and re-experiencing symptoms on the Clinician-Administered PTSD Scale (CAPS-B) in an overlapping aspect (purple) of the ventromedial prefrontal cortex, corresponding to BA10.

## 4. Discussion

In a study of combat-exposed male veterans, we simultaneously investigated relationships between vmPFC responses to safety vs. threat, resting high-frequency pulse rate variability, and PTSD re-experiencing symptoms. Three key findings emerged from this investigation that shed light on potential brain-body mechanisms of PTSD.

First, in veterans with elevated symptoms of PTSD (CAPS > 20), we identified a specific relationship between greater re-experiencing symptoms of PTSD and reduced PRV, a surrogate measure for HRV that can be obtained in the MRI environment. Relationships between re-experiencing symptoms in PTSD and executive control deficits (Aupperle, Melrose, Stein, & Paulus, 2012; Bomyea, Amir, & Lang, 2012; Vasterling, Brailey, Constans, & Sutker, 1998) suggest that reduced inhibitory control compromises the ability to prevent unwanted traumatic memories from rising into awareness in PTSD. Further, it has been proposed that HRV indexes one’s ability to implement flexible regulatory control in the face of distractors or challenges to one’s well-being (Thayer et al., 2012), and HRV has been linked to difficulties suppressing unwanted memories in healthy college students (Gillie, Vasey, & Thayer, 2014). Gillie & Thayer (2014) proposed that reduced HRV contributes to the inability to control unwanted memories or thoughts related to trauma and as such represents a physiological mechanism of re-experiencing symptoms, such as flashbacks and nightmares. Our results provide empirical support for this theoretical model, although replication of these results in a larger sample that also incorporates behavioral indices of cognitive control is an important next step.

The second key finding was that reduced PRV was associated with reduced vmPFC recruitment under conditions of safety vs. threat. Convergent clinical observations and neuroscientific evidence suggests that PTSD is not associated with exaggerated responses to threat *per se*, but rather contextually inappropriate, inflexible, and overgeneralized threat responding (Garfinkel et al., 2014; Kaczkurkin et al., 2017; Levy-Gigi, Richter-Levin, Szabo, & Keri, 2015; Morey et al., 2015). An inability of the vmPFC to differentially respond to threat vs. safety, and corresponding autonomic inflexibility reflected in reduced PRV, may contribute to elevated threat responding in an objectively safe context. The region of vmPFC identified here corresponds to frontopolar cortex or medial BA10, a region that has expanded considerably in humans relative to non-human primates and which is theorized to be important for shifting attentional or executive resources away from current goals to other potential goals in the environment (Mansouri, Koechlin, Rosa, & Buckley, 2017). This perspective on the broad function of BA10 is not inconsistent with the flexible, context-specific, inhibitory control function ascribed by Thayer and colleagues to HRV (Thayer et al., 2012). Notably, a previous study identified a positive correlation between HRV and activation in a similar vmPFC region during self-control challenges (Maier & Hare, 2017), a finding consistent with the current findings in light of the above framework linking re-experiencing symptoms to deficient cognitive control.

Third, this relationship with reduced HF-PRV was spatially overlapping with a BA10 region in which we previously identified a relationship with re-experiencing symptoms in these same participants. This anatomical overlap provides tentative support for the idea that observations of reduced vmPFC function and decreased HRV/PRV in PTSD may be mechanistically related, although stronger support for this hypothesis would require alternative designs. For example, it would be informative to test whether the effectiveness of somatic interventions targeting PTSD symptoms – for example, aerobic exercise (Fetzner & Asmundson, 2015), meditation (Polusny et al., 2015), yoga (Gallegos et al., 2017), or HRV feedback training (Tan, Dao, Farmer, Sutherland, & Gevirtz, 2011) – is predicted by engagement of this hypothesized target mechanism, i.e., normalized vmPFC function and corresponding increases in HRV.

One question raised by our results is why control participants with the lowest HF-PRV levels had few or no re-experiencing symptoms. HF-PRV was, on average, no different between CEC and PTSS groups and was related to re-experiencing symptoms only in the PTSS group (**Figure 2a**). These observations underscore that although reduced HRV/PRV may contribute to (or be exacerbated by) PTSD intrusions, it is by no means a sufficient condition. Control participants with low levels of HRV/PRV may possess compensatory mechanisms that enhance resilience against psychopathology. For example, these individuals may be better able to leverage explicit, cognitive emotion regulation resources – supported by dorsal and lateral PFC (Buhle et al., 2013) – to compensate for deficient autonomic regulatory control. Alternatively, they may exhibit enhanced hippocampal function that allows them to better differentiate between safe and threatening contexts (Anacker & Hen, 2017). The concurrent investigation in traumatized individuals of HRV/PRV and neuroimaging investigations of explicit emotion regulation or hippocampal-dependent tasks (e.g., pattern separation behavior) would allow for a test of these speculative hypotheses.

One important limitation of these results is that PRV and HRV, while measuring the same underlying signal, are not fully equivalent measures. We assessed PRV using pulse oximetry due to the difficulty of collecting electrocardiography data in the MRI environment. Although PRV and HRV are highly correlated, particularly at rest (Hayano et al., 2005; Schäfer & Vagedes, 2013), the two measures rely on distinct physiological readouts of cardiac function and PRV may be less accurate for the measurement of high-frequency variability in particular (Wong et al., 2012). Much of the extant literature on PTSD and cardiac autonomic control assesses HRV and not PRV, (although some published “HRV” studies in fact utilize photoplethysmography; e.g., Minassian et al., 2015), and caution is warranted in generalizing the current results to the HRV literature until PRV is better established as an index of regulatory control and psychopathology. An additional important limitation is that we studied a relatively homogeneous sample (male OEF/OIF veterans who experienced combat trauma), which while reducing potential sources of variability also limits generalizability of findings to female veterans or individuals exposed to non-combat trauma.

In summary, we have presented evidence that reduced functional activation in spatially overlapping voxels of the vmPFC is associated with both reduced HF-PRV and increased re-experiencing symptoms of PTSD. These data provide preliminary support for a mechanistic link between previous observations of reduced HRV and impaired vmPFC function in PTSD. More broadly, these findings underscore the potential of research on reciprocal brain-body relationships that may enhance our understanding of the pathophysiology of PTSD and suggest novel treatment targets for somatically focused therapeutic approaches.

## Author Notes

The authors thank the participants for their military service and their involvement in this study, as well as the Wisconsin National Guard, the Madison Veterans’ Center, the Madison VA Hospital, and other veterans’ community organizations for their assistance in recruitment. The authors thank Kate Rifken, Andrea Hayes, Emma Seppala, Michael Anderle, Lisa Angelos, Isa Dolski, Ron Fisher, and Nate Vack for their help with study planning, execution, and analysis, and Jared Martin for thoughtful feedback on this manuscript.

This work was supported by the Dana Foundation to JBN; the University of Wisconsin Institute for Clinical and Translational Research to Emma Seppala; and a core grant to the Waisman Center from the National Institute of Child Health and Human Development (P30-HD003352). DWG was supported by a Graduate Research Fellowship from the National Science Foundation.

Portions of this work were previously presented at the 72^nd^ Annual Scientific Convention of the Society of Biological Psychiatry, San Diego, May 18, 2017.

Dr. Davidson is the founder and president, and serves on the board of directors, for the non-profit organization Healthy Minds Innovations, Inc. The other authors report no conflicts of interest or financial disclosures.

